# Scalable Integration of Multiomic Single Cell Data Using Generative Adversarial Networks

**DOI:** 10.1101/2023.06.26.546547

**Authors:** Valentina Giansanti, Francesca Giannese, Oronza A. Botrugno, Giorgia Gandolfi, Chiara Balestrieri, Marco Antoniotti, Giovanni Tonon, Davide Cittaro

## Abstract

Single cell profiling has become a common practice to investigate the complexity of tissues, organs and organisms. Recent technological advances are expanding our capabilities to profile various molecular layers beyond the transcriptome such as, but not limited to, the genome, the epigenome and the proteome. Depending on the experimental procedure, these data can be obtained from separate assays or from the very same cells. Despite development of computational methods for data integration is an active research field, most of the available strategies have been devised for the joint analysis of two modalities and cannot accommodate a high number of them.

To solve this problem, we here propose a multiomic data integration framework based on Wasserstein Generative Adversarial Networks (MOWGAN) suitable for the analysis of paired or unpaired data with high number of modalities (>2). At the core of our strategy is a single network trained on all modalities together, limiting the computational burden when many molecular layers are evaluated. Source code of our framework is available at https://github.com/vgiansanti/MOWGAN.

## Introduction

Availability of single-cell sequencing technologies made easier the understanding of complex organization of tissues and organs and the interplay among different types of cells. Here, cell properties can be characterized at different layers, in terms of transcriptome (1–4), genome (5–7) epigenome (8–10) and proteome (11,12). Depending on the experimental procedure, these data are available from the same cells or from separate assays (13). Multiomic data from the same cells are currently limited to up to 2/3 modalities (or layers) but the prospective is to have a higher rate of co-observation (14,15), as long as additional assays do not compromise profiling of other layers, keeping the cell intact.

Proper integration of multiomic data is one of the grand challenges in single-cell data analysis (16). Integration is considered a difficult task as it requires specific computational models supporting multidimensional data, it is based on generally unknown models of mutual dependence and causal relationships among modalities, and it includes all the analysis challenges that are peculiar to each single modality (*e.g.*, sampling bias, high sparsity).

Data integration links different data sources to derive a more comprehensive and biologically meaningful description of the object under analysis. This task can be addressed in three different ways depending on the type of available anchors used to combine multiple data sources (17): we refer to *horizontal integration* when integration is performed between different datasets representing the same data type (*i.e.*, multiple scRNA-seq dataset) collected from multiple samples at different locations or time points. V*ertical integration* is performed when more modalities are assayed from the same cell, for example epigenome and transcriptome (10,18,19) transcriptome and proteins (20), genome and transcriptome (21) or any other combination (22–25). Vertical integration is challenging both from an experimental and an interpretative point of view. Finally, we refer to *diagonal integration* if both cells and features are different for all datasets. This last scenario may be considered the hardest to solve and yet it is possibly the most common, given the pace at which single-cell datasets are produced (26).

While horizontal integration can be treated as a problem of batch correction (27), vertical and diagonal concern with the multimodality integration field. To the best of our knowledge, a fully integrated and generalizable way to analyse such data is still missing. The main solutions proposed so far refer to *Manifold Alignment* (MA) applications and *Deep Learning* (DL) (28–42). Both approaches share the final goal of representing multiple feature sets in a common manifold embedding. MA methods try to find a common latent space (manifold) to describe the data, while DL methods develop networks specific for the single omic and work on their embeddings, (*i.e.*, learned low-dimensional representations of the given data).

In order to produce their results, both MA and DL methods must assume some constraints. Restrictive assumptions are generally applied on the data, like correspondences between the features (*e.g.*, converting chromatin accessibility data into gene activity scores (43)) and/or between cells, or assumptions on the data distribution. These requirements are difficult to fulfil and to generalize, making them unfit for datasets where no prior knowledge is available. Furthermore, they are typically considered to address the integration of only two molecular layers (*e.g.*, RNA and ATAC), so scaling to three or more modalities could be impractical.

In addition to technical limitations and experimental costs, many large datasets have been made available for single omics only (44–46). This urges the need for the development of an approach to integrate paired or unpaired data which is also free from assumptions and flexible with respect to the number of omics available. To address these issues, we propose MOWGAN, a DL framework for the analysis of Multi-Omics paired or unpaired data based on Wasserstein Generative Adversarial Networks. Our approach is designed to accommodate any kind and number of single-cell assays without priors on the relationships among the inputs. MOWGAN learns the structure of single assays and infers the optimal couplings between pairs of assays. In doing so, MOWGAN generates synthetic multiomic datasets that can be used to transfer information among the measured assays by bridging (47). We benchmarked MOWGAN with existing methods for multimodal single-assays data integration on various single cell datasets and show how it could be used to unveil hidden biology. We show that MOWGAN generates much more reliable results when looking at the cell type annotation shared between the layers.

A python implementation of MOWGAN can be found at https://github.com/vgiansanti/MOWGAN.

## Results

### Overview of MOWGAN

The core component of the framework is a Wasserstein Generative Adversarial Network (WGAN) with gradient penalty (WGAN-GP). A WGAN-GP is a generative adversarial network that uses the Wasserstein (or Earth-Mover) loss function and a gradient penalty to achieve Lipschitz continuity (48,49). Like any other GAN, the WGAN-GP is composed of two subnetworks, called *generator* and *critic*. MOWGAN’s *generator* outputs a synthetic dataset where cell pairing is introduced across multiple modalities.

MOWGAN’s inputs are molecular layers embedded into a feature space having the same dimensionality *C* (Figure 1). In principle any dimensionality reduction technique could be used (*e.g.*, PCA, SVD…). To capture local topology within each dataset, cells in each embedding are sorted by the first component of its Laplacian Eigenmap (LE). This step maximises the likelihood of having similar cells close to each other within each layer. Training of the WGAN-GP is performed in mini-batches, iteratively sampling random subsets of cells from the whole sorted dataset. First, a mini-batch is selected from one modality (usually RNA) and a Bayesian ridge regressor *R* is trained on the mini-batch embedding and the corresponding eigenvectors from the LE. Next *n* mini-batches (default *n* = 50) are selected from the second modality. We apply *R* on all mini-batches and their eigenvectors. The resulting scores are used to sort mini-batches and select the batch in the second modality (*e.g.*, ATAC) with local properties more similar to the first, therefore maximising the probability to select cells drawn from same cell types. This procedure is repeated for all modalities, always using one layer (*e.g.*, RNA) as reference. In preliminary implementations of MOWGAN, without LE sorting and pre-matching, we noticed the WGAN-GP could only capture global topology of the data, resulting in random mixing of cell types (Figure S1).

**Figure 1:**
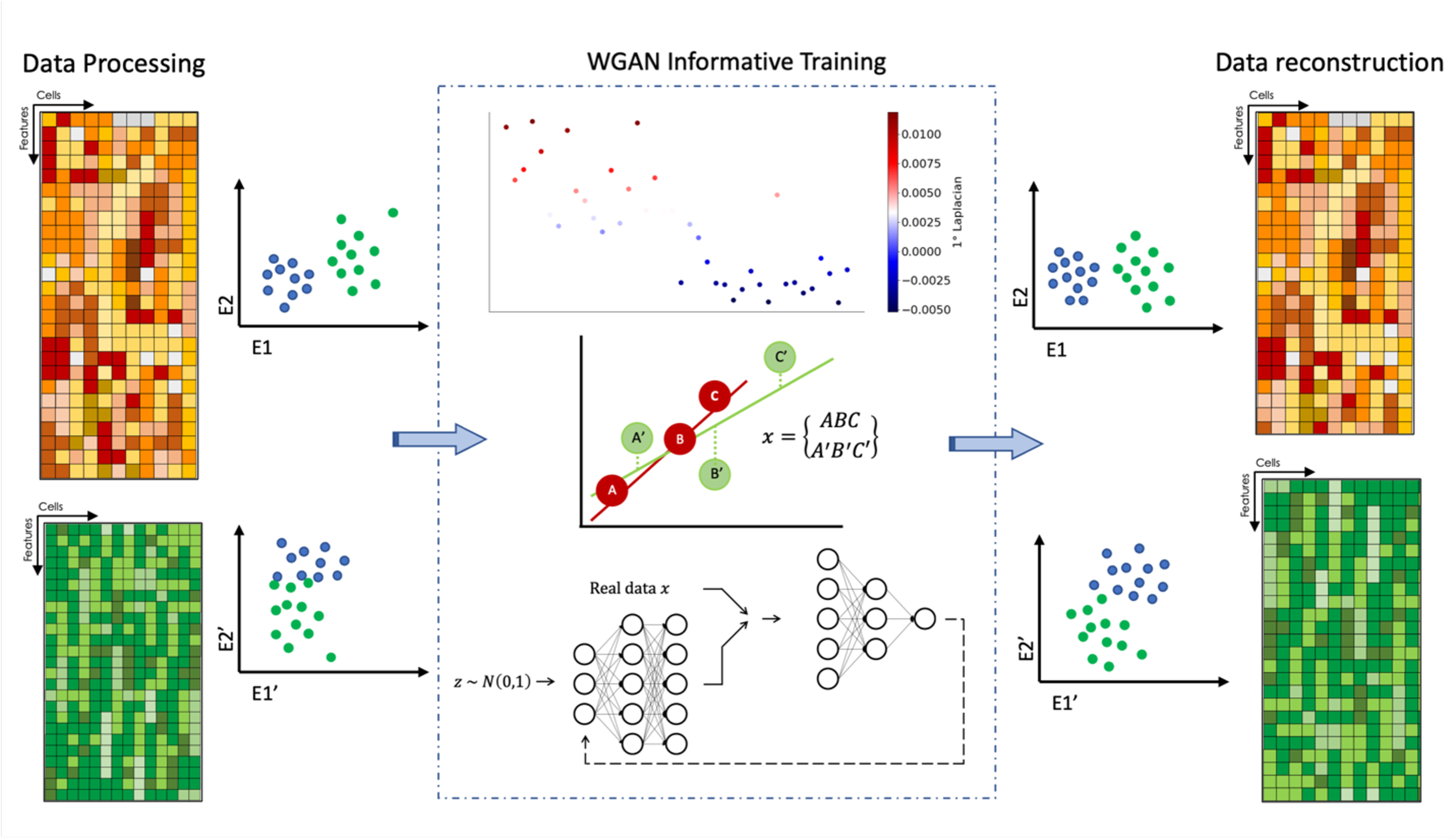
Overview of MOWGAN. MOWGAN is a framework for the synthetic generation of multi-omics data. Given unpaired single cell data (*e.g.*, scRNA-seq and scATAC-seq from separate assays), MOWGAN learns both the specificity and the relationship between modalities, returning a new, paired, dataset. MOWGAN takes the data embeddings as input (step 1), sorts the samples according to the spectral embedding (step 2), and imputes the association between cells of different modalities through a Bayesian Ridge Regressor (step 3). Imputed pairs are used to train a WGAN-GP (step 4). The final output is a single cell multi-omics dataset including embeddings generated by the WGAN-GP and feature count matrices reconstructed using kNN regression (step 5).

The mini-batches are combined to form a tensor (*N*, *M*, *C*), where *N* is the number of cells in a mini-batch (default *N* = 256), *M* is the number of modalities evaluated (2 in case of just the RNA and ATAC layers), and *C* is the rank of the input embeddings. In any case, *C* is set at most to the lowest matrix rank across all modalities.

Semi-supervised training is implemented to include prior information about the dataset. This allows to model, for example, batch-specific models that are eventually merged. Lastly, MOWGAN can explicitly include paired data during training.

We implemented MOWGAN in Python using Tensorflow libraries to support of GPU-enabled hardware and scale to large datasets.

### MOWGAN preserves data topology and biological information

We prototyped MOWGAN on public scRNA and scATAC data of peripheral blood mononuclear cell (PBMC) for which paired and unpaired experiments were available (10x Genomics datasets). The paired dataset (PBMC_1) is a true multimodal experiment, with transcriptome and epigenome data from the same cells; the unpaired dataset (PBMC_2) includes data from two separate experiments, one for each modality.

We evaluated MOWGAN output in terms of shared information between layers (Adjusted Mutual Information of cell clusters, AMI), integrability (Local Inverse Simpson’s Index, LISI (50)) and ability to recapitulate biological information within the same modality. Experiments were performed using either PBMC_1 or MOWGAN as data bridge to transfer labels from RNA to ATAC modality.

First, we identified a set of cell clusters *T* (“truth”) in each modality, we used data bridges (either the PBMC_1 dataset or synthetic pairings generated by MOWGAN) to transfer labels from one modality to another. We denote the set of transferred cell clusters as *P* (“prediction”). Labels are propagated from the first modality *D*_1_ to MOWGAN and then from MOWGAN to the second modality *D*_2_ using label transfer properties of a Stochastic Block Model (51). As every cell in *D*_1_ and *D*_2_ will be annotated by *T* and *P*, we could calculate mutual information. When measured on PBMC_1 data using direct transfer by cell identity, mutual information was computed comparing *T* annotations of RNA and ATAC and found below 1 (AMI = 0.691), indicating that different modalities encode slightly different cell properties.

We obtained lower mutual information values when cell clusters were transferred using MOWGAN, which was expected (AMI_RNA_ = 0.662, AMI_ATAC_ = 0.594 for PBMC_1, AMI_RNA_ = 0.411, AMI_ATAC_ = 0.448 for PBMC_2). Lower values for PBMC_2 were also expected, as the dataset is composed by two unpaired experiments which were acquired, analyzed and released over a long extent.

Data integrability was measured by LISI, a measure of the amount of data mixing in the cell neighborhood. If two datasets (*A,B*) are well integrated, we may expect the neighbors of every cell in *A* being well represented in *A* and *B* datasets, and *vice versa*. When integration fails, most of the neighbors of every cell in *A* (*B*) will be from dataset *A* (*B*) only. LISI is calculated within the same modality (*D*_1_ or *D*_2_ with MOWGAN) and it should be intended as a global measure of similarity of two datasets. It varies from 1 (perfect separation) to the number of modalities *M* (perfect mixing).

Baseline measures of integrability, evaluated on PBMC_1 and PBMC_2, were dependent on the modality (LISI_RNA_ = 1.2, LISI_ATAC_=1.35). Data generated by MOWGAN had higher LISI values (LISI_RNA_ = 1.636, LISI_ATAC_ 1.641 for PBMC_1 and LISI_RNA_ = 1.354, LISI_ATAC_ = 1.438 for PBMC_2), indicating that it is able to compute synthetic data that resemble the input.

Lastly, we evaluated the ability to transfer biological information in terms of power of identification of cell types. We identified sets of cell type markers in each modality of PBMC_1. We then calculated, for each cell type, the fold change of each feature in MOWGAN data and PBMC_2, the latter to evaluate baseline performances, and performed preranked GSEA (52) against the markers. While the WGAN-GP itself does not generate feature values, MOWGAN imputes them using kNN-regression (see Methods). We evaluated the performance as average rank *K* of NES values on the diagonal in the confusion matrix, so that *K* = 1 if all enrichments maximally match the correct cell type and *K* = 0 if the correct cell type has the lowest possible enrichments (Figure 2).

**Figure 2:**
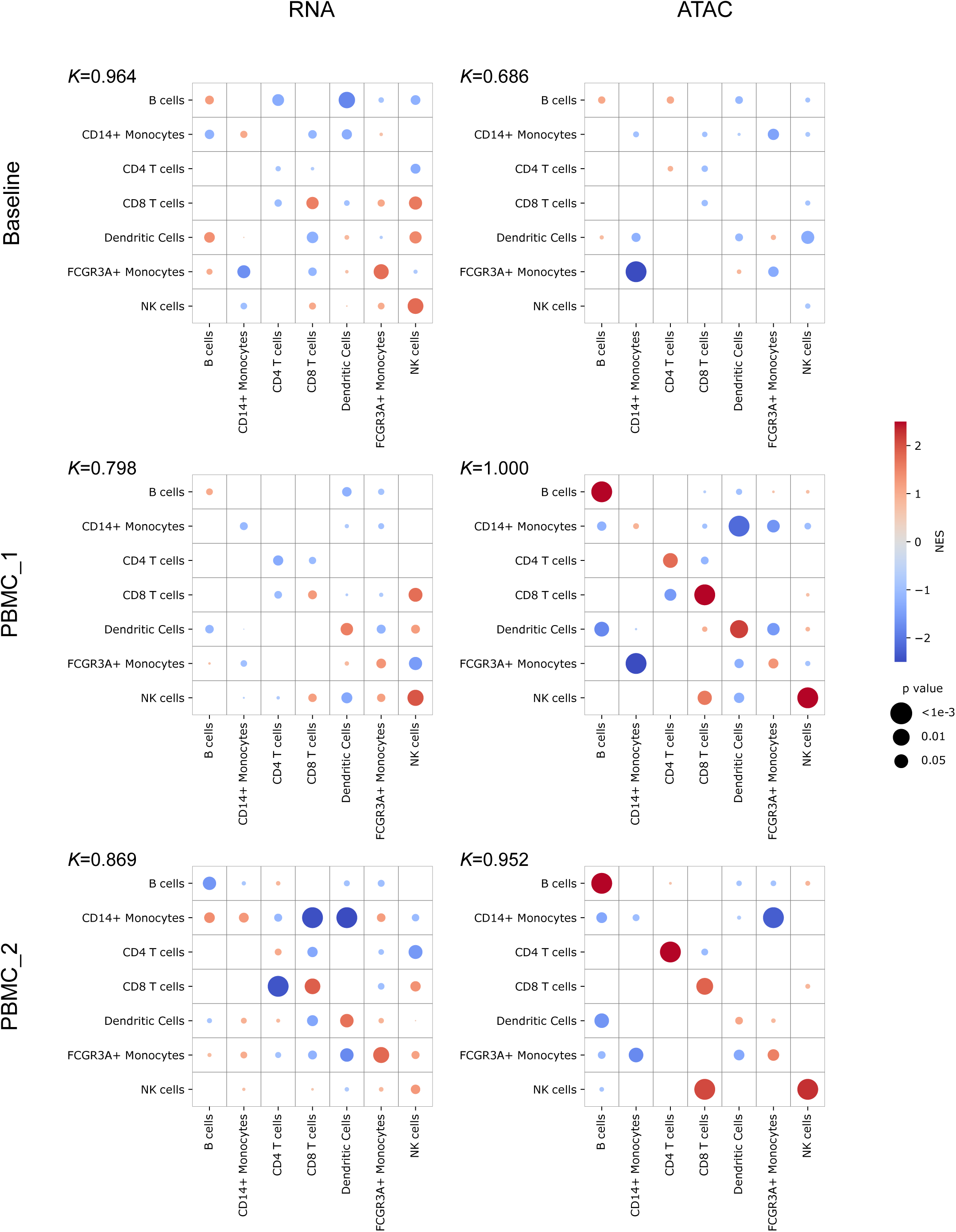
Enrichment analysis of biologically relevant features. Within each matrix, each column represents the result of preranked GSEA of reconstructed feature values for different cell types, compared to features identified as cell type markers in the input data. In the baseline analysis (top row), any single modality (RNA or ATAC) of PBMC_2 is compared to the same modality of PBMC_1. MOWGAN outputs for PBMC_1 (center row) or PBMC_2 (bottom row) are compared to the relative modality of input data. The left column reports results for scRNA-seq, the right column reports results for scATAC-atac. Colour is proportional to NES value, dot size is proportional to the p-value of the enrichment. We report an accuracy metric (*K*) which is the average rank of NES values on the diagonal.

The baseline performance, evaluated with PBMC_2, indicates that different datasets, while derived with different protocols, share a large portion of explained biology (*K*_RNA_ = 0.964, *K*_ATAC_ = 0.686). Synthetic data generated with MOWGAN share much of the biological information with the respective input data, performance was high both for PBMC_1 (*K*_RNA_ = 0.800, *K*_ATAC_ = 1.000) and PBMC_2 (*K*_RNA_ = 0.869, *K*_ATAC_ = 0.952). We noticed that *K*_ATAC_ for MOWGAN was higher than baseline, we ascribe this result to the intrinsic denoising properties of the generative model and the kNN regressor.

In all, these results indicate that MOWGAN can induce appropriate pairings in synthetic data which, in turn, can be used to perform bridge integration of multiple modalities.

### Comparison with other tools

We benchmarked MOWGAN against four tools that share similar design principles: Pamona (30), SCOT (32), COBOLT (35) and scMMGAN (29). Pamona and SCOT use Optimal Transport theory to project each modality on a single latent embedding. The new embeddings can be processed by standard workflow to identify cell clusters. scMMGAN projects data of each modality to the embedding defined on the other modality, so that two modalities can be aligned in a single space. Similarly to Pamona and SCOT, the resulting embedding can be used to identify cell clusters. COBOLT, instead, projects unpaired data on the embedding obtained from paired data, to perform bridge integration.

For each tool, we evaluated the consistency between cell types and cell clusters derived from the generated embeddings, both in PBMC_1 and PBMC_2 datasets. Cell types were annotated once on both datasets, to be consistent across all experiments. In addition to AMI between *P* and *T* cell clusters, we measured Completeness (*i.e.*, all cells of any cell type are clustered together), Homogeneity (*i.e.*, all cells of any cluster belong to the same cell type), and their harmonic mean (V-measure). In general, we observed better performances in PBMC_1 compared to PBMC_2 and on the RNA layer compared to the ATAC layer, without any tool outperforming the others (Figure 3A, Table S1).

**Figure 3:**
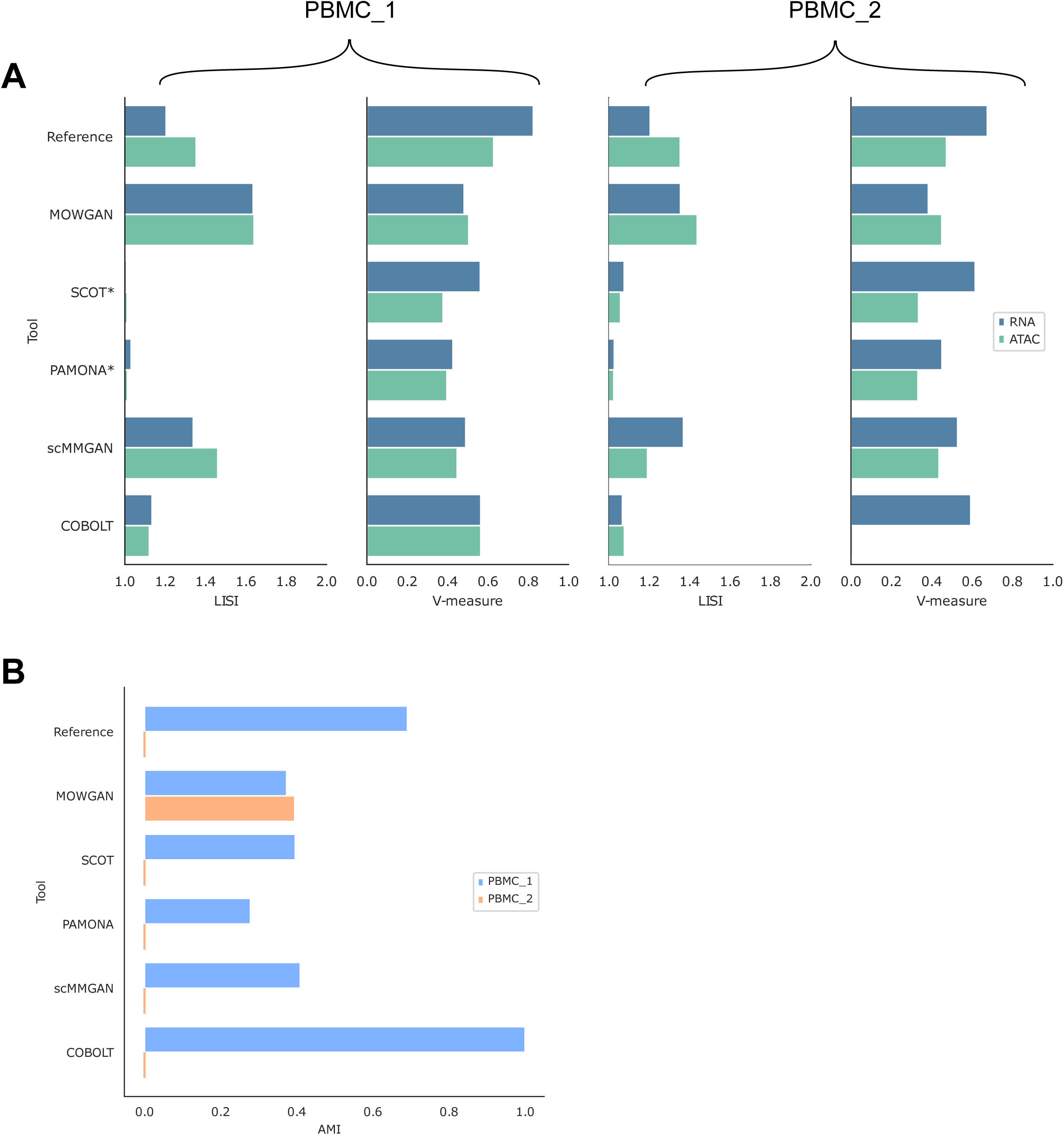
Comparison with other tools for multimodal single-cell integration. (A) The histograms display LISI and V-measure scores. V-measure is obtained comparing annotated cell types and cell clusters identified in each modality after data integration. Two groups of histograms are reported for PBMC_1 and PBMC_2 datasets separately. Both SCOT and Pamona have been marked by an asterisk since the LISI value has no meaningful interpretation, due to the fact both generate a new embedding in a coordinate system different from the respective input. (B) Histogram displaying the AMI values between cell clusters identified in scRNA-seq and scATAC-seq data after data integration.

More in detail: SCOT had the best V-measures for RNA (V_PBMC_1_ = 0.562, V_PBMC_2_ = 0.616) and the worst for ATAC (V_PBMC_1_ = 0.378, V_PBMC_2_ = 0.335), whereas MOWGAN displayed a similar performance both on RNA (V_PBMC_1_ = 0.482, V_PBMC_2_ = 0.383) and ATAC (V_PBMC_1_ = 0.505, V_PBMC_2_ = 0.450). MOWGAN also performed similar to scMMGAN, with which shares some design principles as both rely on deep generative models. Of note, COBOLT failed in integrating ATAC data in PBMC_2 (V_PBMC_2_ = 0.009) and its high performance in PBMC_1 was probably due to the fact it explicitly uses cell identities to integrate two modalities.

Measures of mutual information reveal that all tools share similar performance on PBMC_1 dataset (average AMI_PBMC_1_ = 0.364) but MOWGAN was the only tool sustaining comparable performance on PBMC_2 (AMI_PBMC_2_ = 0.395), while the remaining tools had AMI values close to 0 (Figure 3B, Table S1).

Integrability measured by LISI showed comparable performance of MOWGAN and scMMGAN, again possibly due to the fact both are based on GAN. We report low LISI values for SCOT and Pamona, however they should not be considered in the comparison as both tools generate an embedding in a new reference system that is necessarily different from the input embeddings.

Last, we noticed some tools induce spurious cell type associations (Figure S2). For example, SCOT switched B cells with NK/CD8 T cells in PBMC_1 and Monocytes with T cells in PMBC_2. Similarly, scMMGAN suffers of several cell type switches in all the tested datasets. As previously noticed, COBOLT shuffled cell types in the ATAC layer in PBMC_2.

In all, these results indicate that all tools but COBOLT preserve the local topology of the input data as clustering properties are conserved in the data transformations. In this regard, no tool outperforms the others, suggesting an upper bound for integration tasks. However, most tools, except for MOWGAN, did not guarantee the correct cell type associations, a further evidence of the ability of MOWGAN to generate good quality and meaningful data pairings.

### MOWGAN extends the integration to high number of modalities

The major advantage of MOWGAN is that it can integrate more than two molecular layers without modifying the model architecture. To test this feature we took advantage of the availability of different single cell profiles for PBMC; in particular we attempted the integration of PBMC_1 dataset with H3K27me3 and Antibody Derived Tags (ADT) data obtained with scCUT&TAG-pro (53) on PBMC, to achieve a four-layers integration. It’s worth underline that the pairings between ADT and H3K27me3 and between RNA and ATAC were not exploited in this experiment. We observed high integrability between original and synthetic data generated with MOWGAN, with the lowest performance detected for H3K27me3 (LISI_RNA_ = 1.67, LISI_ATAC_ = 1.67, LISI_ADT_ = 1.75, LISI_H3K27me3_ = 1.38).

We used MOWGAN to bridge cell type annotations from RNA to all the remaining modalities, then compared with the given cell type (Figure 4). We found similar results for ATAC and ADT, both in terms of accuracy (> 0.8) and V-measure (> 0.6), whereas H3K27me3 performed worst, suggesting RNA, ATAC and ADT share similar resolution power with respect to cell types. On the other hand, H3K27me3 clearly separated myeloid and lymphoid lineages with lower resolution power in the latter. An analogous behavior of H3K27me3 has been previously described (54).

**Figure 4:**
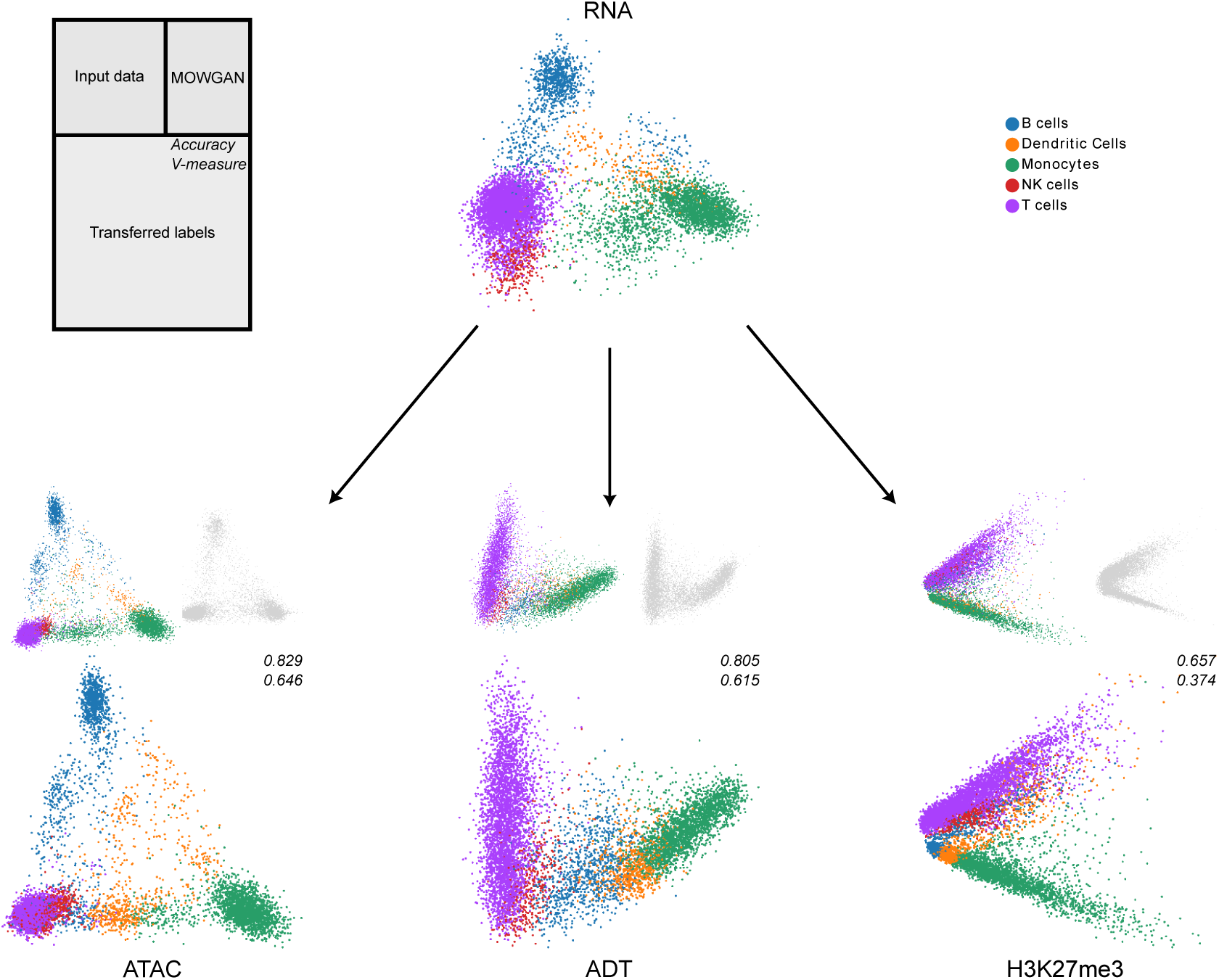
Integration of multiple single cell modalities. We report the low dimensional embeddings (PCA) of multiple single cell profiles of PBMC. After data integration, we used MOWGAN to bridge cell type annotations from RNA to ATAC, ADT and H3K27me3 layers. For each modality receiving the annotation we report three embeddings: the input data colored by cell type (top left), the embedding of data generated by MOWGAN (top right) and the input data colored by the transferred cell type (bottom). We report values of accuracy and V-measure for each transfer as well.

Of note, we found the performance of inverse labeling, from any modality to RNA, equally high, if not better (Table S2). In all, these results show how MOWGAN can effectively compute synthetic pairings for multiple modalities, overcoming a limitation that is common to many available tools.

### Semisupervised training improves results and unveils hidden cell populations in colorectal cancer organoids

An ideal tool to integrate unpaired data would be able to work in unsupervised manner, that is it would learn the whole data structure from data themselves. That requires the algorithm to model several confounders including technical noise due to different technologies (13,55), biological noise (56,57) and batch effects (27). For this reason, we tested the ability of MOWGAN to work with multiple samples of unpaired data with or without specifying the sample identity in the training step.

We profiled three patient-derived organoids (PDO) of liver metastatic colorectal cancer (CRC) by scRNA-seq and scGET-seq (25) (Table S3). Cells for each PDO were split in two aliquots before single cell preparation, so that data are paired at sample level (*i.e.*, single cells represent the same populations) but not at modality level. Analysis of copy number alterations at single cell level revealed that all PDOs are genetically uniform and essentially monoclonal (Figure S3A). These findings were previously confirmed for CRC6 and CRC17 by bulk exome sequencing (25).

To verify the effectiveness of semisupervised training, we trained one MOWGAN model without specifying the organoid of origin (NB) and a model that instead included such information (B). In practice, to obtain model B, we trained three separated models, one per organoid, sampling batches from the whole dataset, and eventually concatenating the outputs.

We checked the average concordance between the actual organoid labels and the one we obtained transferring it from scRNA-seq to scGET-seq and *vice versa* using MOWGAN as bridge. In model NB the performance was low in terms of accuracy (ACC_RNA_ = 0.111, ACC_GET_ = 0.353) and increased in model B (ACC_RNA_ = 0.982, ACC_GET_ = 0.668).

Synthetic data were generally integrable with the original ones, with minor differences between model B (LISI_RNA_ = 1.55, LISI_GET_ = 1.33) and NB (LISI_RNA_ = 1.42, LISI_GET_ = 1.25), in line with previous observations on PBMC datasets. In all, we conclude that semisupervised learning improves the quality of the generated data and their usability for the integration.

Having set the model B is better, we performed further analysis to unveil the biological features of PDOs. We identified cell states using the multimodal capabilities of schist (51), which relies on a Stochastic Block Model with overlap. Two kNN graphs, including MOWGAN generated cells, are computed for each modality and merged specifying an edge label depending on the modality, then a Nested Stochastic Block Model (NSBM) with edge covariates is inferred. Of note, MOWGAN output is represented in the graph as a set of nodes having both types of edges, hence providing the link between modalities. The resulting groups are consistent across modalities and can be used to analyze phenotypes from the transcriptomic and the epigenetic standpoint. We computed a unified embedding using optimal transport framework to visualize data annotations in consistent way (Figure 5A-C).

**Figure 5:**
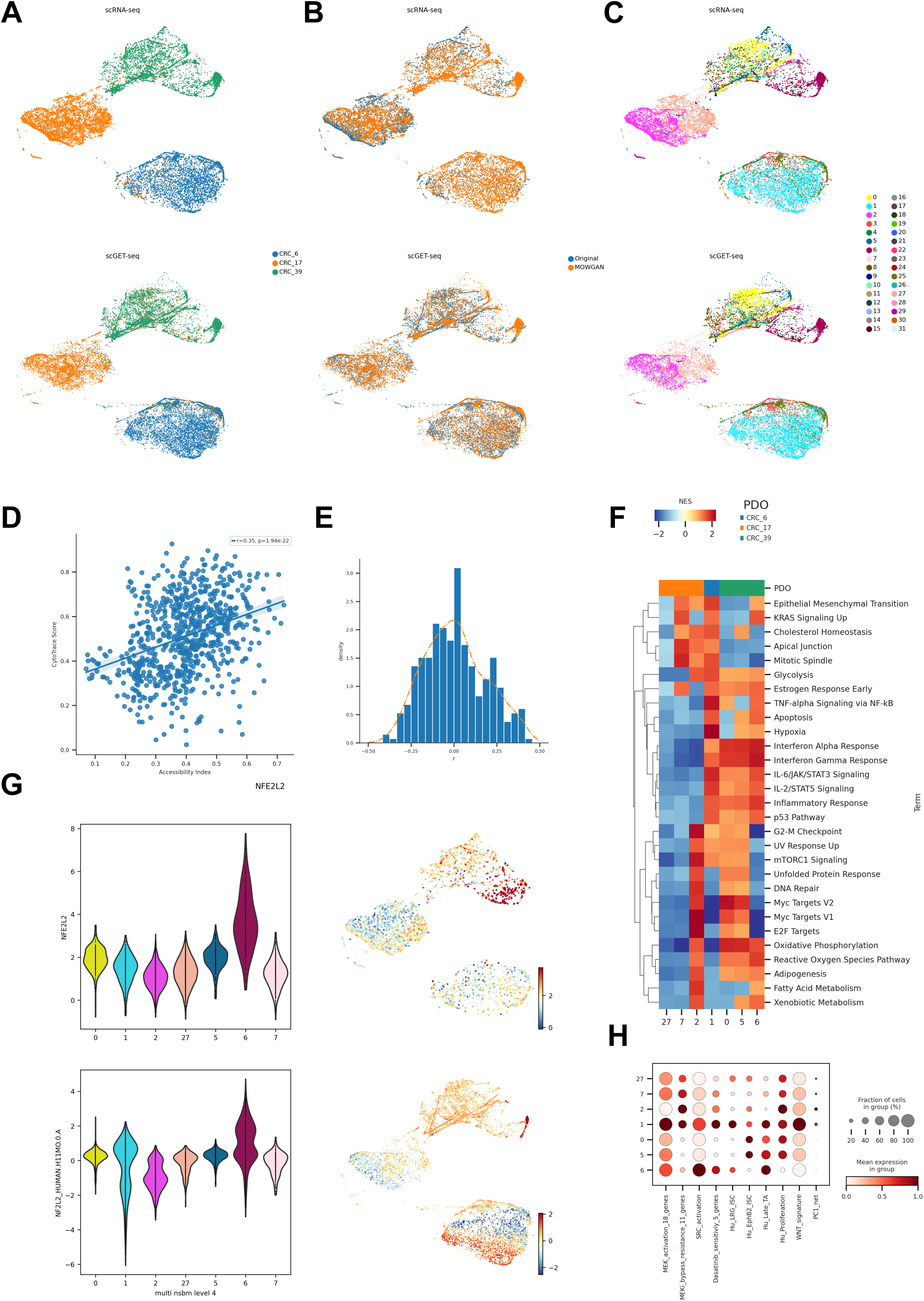
Analysis of CRC PDOs. (A) Unified UMAP embeddings of the scRNA-seq (top) and scGET-seq data (bottom) where single cells have been colored by PDO of origin. (B) Unified UMAP embedding as in A, cells have been colored according to their data source, whether it is original input data or synthetic data generated by MOWGAN. (C) Same embedding as in A and B, cells have been colored by the level 4 of the NSBM hierarchy inferred on multimodal data. (D) Scatterplot representing the correlation between Chromatin Accessibility Score, calculated on scGET-seq data, and CytoTrace score, calculated on scRNA-seq data. Each point represents the median value of a score in cell groups at level 0 of the NSBM hierarchy. (E) Distribution of the correlation values between Transcription Factor activities, evaluated on scRNA-seq data, and Total Binding Affinity, evaluated on scGET-seq data. The orange line traces the smoothed profile of the histogram (F) Gene Set Enrichment Analysis of most represented cell groups after integration. The heatmap displays the Net Enrichment Score of preranked GSEA, using Hallmark gene sets. (G) NFE2L2 profiles in integrated data. We report the distribution of TF activity and TBA score of NFE2L2 in scRNA-seq (top) and scGET-seq (bottom) for the most represented cell groups. Next to each violin plot we report the relative unified UMAP embedding, colored by the same score. (H) Profile of gene signatures described for drug resistance phenotypes in CRC. The dotplot reports the score of each signature in the most represented cell groups found in integrated data.

Since the rate of transcription depends, among other factors, on genome accessibility, we calculated the CytoTrace score (58) from scRNA-seq and a chromtain accessibility score (59) from scGET-seq. The correlation between the two scores, averaged on the deepest level of NSBM hierarchy, was positive (Figure 5D, *r* = 0.35, *p* = 1.94e-22). In addition, we found a positive correlation between accessibility and CRIS-B signature for CRC (Figure S3B, *r* = 0.18, *p* = 4.054e-7). We expected positive correlations in case of appropriate pairing, since CRIS-B subtype has been described associated to EMT phenotype (60), as well as higher CytoTrace scores and chromatin accessibility (61,62) from scGET-seq.

We then evaluated the activity of various transcription factors from gene expression data, using decoupler (63) and a cell-level measure derived from Total Binding Affinity (TBA, (64)), computed from chromatin accessibility (see Methods). Correlation coefficients of the two measures were skewed to positive values (Figure 5E, skeweness = 0.251), again supporting the good quality of the integration. Among other transcription factors (Table S4), we found high correlation for NFE2L2 (*r* = 0.39, *p* = 1.742e-28), which regulates stress response (65) and has been recently found involved in CRC resistance to therapy (66). Interestingly, NFE2L2 is strongly associated with only one subpopulation in CRC39 (Figure 5G), indicating the PDOs, while being genetically uniform, display phenotypically diverse cell states.

We focused our attention at level 4 of the NSBM hierarchy as it provided a good trade-off between modularity and detail on the subpopulations within each PDO. We identified 32 clusters, seven of which include more than 200 cells in both modalities. All PDOs, apart from CRC6, were composed of different subpopulations with distinct phenotypes (Figure 5F). We scored multiple expression signatures studied for their relevance in drug resistance (67) and confirmed that both CRC17 and CRC39 included subpopulations which could support a differential response to treatments (Figure 5H, Figure S3C).

Importantly, these cell populations could be hardly identified by using scRNA-seq or scGET-seq data alone. First, we noticed that the NSBM hierarchy of cell clusters identified in single assays did not strictly follow the PDO identities and, in fact, some clusters are shared across different organoids, particularly in the scGET-seq data. Cell groups identified on the integrated data appear to be more consistent with the organoid identity than groups identified on the single modalities (Figure S3D); this behavior is true at any level of the NSBM hierarchies. Consequently, some phenotypes we observed above could not be clearly assigned to specific subpopulations.

In all, these data confirm that MOWGAN is able to produce good quality pairings to perform bridge integration across modalities. Moreover, we show that semisupervised training is an effective approach to operate when multiple samples/batches are available.

## Discussion

Methods to probe molecular phenotypes of single cells at high throughput have been available for more than a decade now. Such methods have been rapidly adopted and are now standard techniques in many fields of biological studies. The largest part rely on analysis of gene expression, for which many technologies are available (68). Methods to investigate single cell epigenomes has also increased in number overtaking scRNA-seq (69). The development of unimodal approaches, including methods to investigate genome sequences (70) or proteomes (71,72), is paralleled by the introduction of multimodal techniques, whose number has also exploded in recent years (14,15,73). Unless innovative chemistry/microfluidic strategies will be introduced, multimodal investigation will be inherently limited to methods that preserve cell or molecule integrity. Moreover, despite the availability of multimodal techniques, their analysis remains challenging (74), moreover they may be less informative of single-assay methods (75). For all these reasons, the development of computational strategies to pair unimodal assays is flourishing.

Along this path, we introduced MOWGAN, a DL-based framework that makes use of Wasserstein Generative Adversarial Networks (WGAN) to generate synthetic multimodal data for bridge integration of different single-cell assays. Given the heterogeneity of available technologies, we designed MOWGAN to work without the need of “biologically principled anchors” across modalities, such as Gene Activity Score for scATAC-seq data (43) or Chromatin Silencing Score for repressive histone marks profiled in scCUT&Tag (54).

Comparison with other tools on the integration of two unpaired modalities (scRNA-seq and scATAC-seq) shows MOWGAN performs similar to the top-level methods, without any tool outperforming the others, suggesting an intrinsic upper bound for this task. The major improvement of MOWGAN over existing tools is instead the possibility to work with more than two modalities. To do so, it aligns all input data to one base modality of choice, typically the transcriptome. Consequently, MOWGAN does not generate a data barycenter, nor it computes an embedding in a latent space. In this regard, MOWGAN could be used in a modular workflow, allowing the adoption of the method of choice to bridge information. We are aware that other tools can operate on high number of modalities, however they require partial overlaps among data (76) or explicit code refactoring (29).

Another important feature of MOWGAN is the possibility to specify experimental pairings, being those cell or sample identities. Semisupervised training of MOWGAN, when possible, reduces the possibility to hallucinate, a well-known issue that affects deep generative models (77), and allows the computation of good-quality and biologically meaningful data.

While it is established that “no method works best for all”, we believe that MOWGAN has sufficient power to generalize over various technologies. We showed that integration of multiple modalities, notwithstanding that data are generated from separate experiments, provides information in a synergistic way, hence revealing hidden patterns. Our findings are particularly relevant in the context of intratumor heterogeneity. In our PDO dataset, all but one samples have been derived from tumors previously exposed to chemotherapy, therefore it’s expected they contain (epigenetic) subclones selected for drug resistance; CRC39, on the other hand, has been derived from a naïve tumor and it would be tempting to speculate on the existence of subclones conferring drug resistance that could eventually expand upon therapy.

## Materials and methods

### WGAN-GP architecture

MOWGAN core component is a WGAN-GP composed of two networks, a *generator* and a *critic*. The *generator* is designed with 3 *convolutional 1D* layers (Conv1D) and 2 *batch normalization* layers (BN). The *critic* is designed with 2 Conv1D layers and a single Dense layer with 1 unit. All Conv1D layers have common parameters: *kernel size* = 2, *stride* = 1, *activation function* = ReLu. The number of *filters* for Conv1D layer changes with the network depth: default number of filters for the *generator* are 512, 128 and 20, for the *critic* 128 and 512.

Subnetwork uses different optimizers: Adam optimizer for the *generator* with *learning rate* = 0.001, *beta_1* = 0.5, *beta_2* = 0.9, *epsilon* = 1e-07; RMSprop optimizer with *learning rate* = 0.0005 in the *critic*.

### Grid search

We applied a grid search policy to test the dependency of MOWGAN to hyperparameters. We tested multiple filters *f* = {8,32,64,128,256,512} in the Conv1D layers, and number of components selected from the embeddings *C* = {5,10,15}. We also set *C* as the number of filters in the *generator*’s hidden layer. A total of 216 models were trained, with training time for single model of approximately 3h every 100,000 epochs. 51 models returned NaNs in the loss function, 1 model produced data collapsed in a single point.

### Feature reconstruction

We fit a kNN-regressor with *k* = 2 for each modality to learn the relationship between the count matrix and the embedding in the input data. The regressor was then applied on the WGAN-GP output to predict the count matrix of the synthetic data.

### PBMC datasets

Public data for PBMC were downloaded from 10x Genomics resources site. PBMC_1 dataset includes Single Cell Multiome ATAC + Gene Expression (https://www.10xgenomics.com/resources/datasets/pbmc-from-a-healthy-donor-granulocytes-removed-through-cell-sorting-10-k-1-standard-2-0-0). PBMC_2 includes Single Cell Gene Gene Expression (https://www.10xgenomics.com/resources/datasets/10k-human-pbmcs-3-ht-v3-1-chromium-x-3-1-high) <u>and Single Cell ATAC</u> (https://www.10xgenomics.com/resources/datasets/10-k-peripheral-blood-mononuclear-cells-pbm-cs-from-a-healthy-donor-1-standard-1-2-0).

Transcriptome data were aligned to hg38 reference genome using STARSolo (78), providing gencode v36 as gene model (79). ATAC data were aligned to hg38 reference genome using bwa (80), we then extracted the counts over 5kb windows spanning the entire genome. For all datasets we calculated PCA, which was used as input for MOWGAN. Cell clusters were identified using the Planted Partition Block Model, implemented in schist.

### PBMC scCUT&TAG-pro

H3K27me3 and ADT data for PBMC profiled with scCUT&Tag-pro (53) were downloaded from https://zenodo.org/record/5504061 (81). R objects were converted to hdf5 and then loaded into scanpy objects. We retained cell type annotation provided with the datasets. Data were normalized and log transformed before computation of the PCA embedding.

### Patient-derived colorectal cancer organoids (PDOs)

Samples from three individuals with liver metastatic gastrointestinal cancers were obtained following written informed consent, in line with protocols approved by the San Raffaele Hospital Institutional Review Board and following procedures in accordance with the Declaration of Helsinki of 1975, as revised in 2000. PDO cultures were established as previously reported (25,82).

scGET-seq was performed as previously described (25,83) on a Chromium platform (10x Genomics) using ‘Chromium Single Cell ATAC Reagent Kit’ V1 chemistry (manual version CG000168 Rev C) and ‘Nuclei Isolation for Single Cell ATAC Sequencing’ (manual version CG000169 Rev B) protocols. The provided ATAC transposition enzyme (10x Tn5; 10x Genomics) was replaced with a sequential combination of Tn5 and TnH functional transposons in the transposition mix assembly step. Specifically, a transposition mix containing 1.5 μl of 1.39 μM Tn5 was incubated for 30 min at 37 °C, then 1.5 μl of 1.39 μM TnH was added for a 1-h incubation. Nuclei suspensions were prepared to get 5,000 nuclei as target nuclei recovery. Final libraries were loaded on a Novaseq6000 platform (Illumina) to obtain 100,000 reads per nucleus, and a custom read 1 primer was added to the standard Illumina mixture (5′-TCGTCGGCAGCGTCTCCGATCT-3′).

scRNA-seq was performed on a Chromium platform (10x Genomics) using ‘Chromium Single Cell 3′ Reagent Kits v3’ kit manual version CG000183 Rev C (10x Genomics). Final libraries were loaded on a Novaseq6000 platform (Illumina) to obtain 50,000 reads per cell.

### Analysis of PBMC datasets

Standard processing was applied to filter and normalized the data: for RNA, cells with less than 200 expressed genes and genes present in less than 10 cells were discarded; for ATAC, cells with more than 30% of captured regions and regions common to more than 80% of cells were selected. For PBMC_1, we further removed all cells that were not shared across the modalities.

We used PCA as embedding scheme for all datasets. Cell clusters were identified using a Planted Partition Block Model provided by schist (51).

Cell type annotation was defined on RNA PBMC_1 by evaluating cell markers, as illustrated in the Scanpy tutorial (https://scanpy-tutorials.readthedocs.io/en/latest/pbmc3k.html). The annotation was directly transferred to ATAC PBMC_1 by cell identity. Annotations were transferred to PBMC_2 using label transfer function provided by schist.

When we tested integration of four modalities, known cell types were converted to the ones provided in (53) according to the following schema:

(CD4 T cells, CD8 T cells)_10x_, (CD4 T, CD8 T, Other T)_scCUT&Tag-pro_: T cells (CD14+ Monocytes, FCGR3A+ Monocytes)_10x_, (Mono, Other) _scCUT&Tag-pro_: Monocytes (B cells)_10x_, (B)_scCUT&Tag-pro_: B cells (NK cells)_10x_, (NK)_scCUT&Tag-pro_: NK cells (Dendritic cells)_10x_, (DC)_scCUT&Tag-pro_: Dendritic cells

### Analysis of PDOs

Sequencing reads for scGET-seq were processed as previously described (83). Briefly, we counted the number of aligned reads in 5kb windows spanning the genome, both for tn5 and tnH enzymes. We used raw counts to fit Zero Inflated Poisson distributions at cell level, which were in turn used to estimate the excess of accessible and compact chromatin in each bin. We removed bins with coverage lower than the 90^th^ percentile and cells with less than 2000 counts. The count matrices were subjected to tensor train decomposition (TTD), used as low-dimensionality representation of the data.

Sequencing reads for scRNA-seq were aligned to hg38 reference genome using STARSolo (78), using GENCODE v36 as gene model (79). We removed cells that a) were identified as doublets by scrublet (84), b) had more than 25% of mitochondrial reads and c) having less than 2000 genes identified. After normalization, identification of highly variable genes, regression of cell cycle and scaling, we obtained low dimensional representation by PCA.

We used 20 components of TTD and PCA as input for MOWGAN with default parameters, computing three different models, one for each organoid, which were then concatenated so that output data have a model label. We used schist to transfer batch identity from input data to MOWGAN output and retained only synthetic data having model label equal to the transferred label.

We used schist (51) to infer multimodal cell clusters (*scs.inference.nested_nsbm_multi*) on the integrated dataset. Once cell clusters have been identified, we used the cell marginals (*i.e.*, the probability of a cell to be assigned to any cluster) as lineage specifications in CellRank (85), so that driver features could be identified in any dataset (*e.g.*, gene drivers for scRNA-seq, region drivers for scGET-seq).

In order to produce a unified UMAP embedding, we first computed separate UMAP for RNA and GET layers, initializing with PAGA positions (86) calculated on RNA using level 4 of the NSBM hierarchy. We then performed Domain Adaptation by unbalanced Optimal Transport based on Sinkhorn algorithm (87) implemented in PythonOT (88), using level 4 of the NSBM hierarchy as anchors for semisupervised fit.

Transcription factor activity was evaluated on scRNA-seq data, excluding MOWGAN synthetic data), using decoupler (63) with default parameters. To calculate the activity of transcription factors in scGET-seq data we first calculated Total Binding Affinity (64) of HOCOMOCO v11 motifs (89) in each genomic bin, using bionumpy (90). We selected the top 200 region drivers for each cluster, having positive correlation and q-value lower than 1E-20, for a total of 1253 regions. We then calculated the motif score *M* as

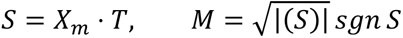

where *X_m_* is the accessibility value estimated on scGET-seq data and *T* is the TBA value for each genomic bin. Once we obtained *M*, we regressed out coverage and scaled the resulting matrix. CRC Intrinsic Subtypes (CRIS) signatures have been collected from CMSCaller package (91).

## Acknowledgements

We would like to thank all members of COSR for the support and fruitful discussions. This work has been supported by the Accelerator Award: A26815 entitled: "Single-cell cancer evolution in the clinic" funded through a partnership between Cancer Research UK and Fondazione Italiana per la Ricerca Sul Cancro (AIRC).

## Author contributions

V.G. and D.C. devised MOWGAN, performed the analysis and wrote the manuscript. V.G. wrote the software. O.A.B. performed the experiments on PDOs. G.G. and C.B. characterized PDOs. F.G. performed single-cell experiments. M.A. and G.T. supervised the work. All authors read the manuscript.

## Data availability

scRNA-seq data for three PDOs have been deposited to ArrayExpress with the following ID: E-MTAB-13123. scGET-seq data for CRC39 have been deposited to ArrayExpress with the following ID: E-MTAB-13126. scGET-seq data for CRC6 and CRC17 were downloaded from ArrayExpress (E-MTAB-10219)

---

Figure S1: Low dimensional embeddings (PCA) of PBMC_1 data generated by WGAN-GP trained with random sampling. The overall distribution of points indicates that the WGAN-GP learns data topology correctly. However, when cell types are transferred across modalities (from RNA to ATAC and vice versa) the cell group information is lost.

Figure S2: Low dimensional representation of PBMC_1 and PBMC_2 datasets generated by different tools used for benchmarking. Each row reports the embedding produced by a tool, colored by cell type. As described in the text, the analysis of these embeddings reveals spurious cell type transfers across modalities for some tools (*i.e.*, SCOT, scMMGAN and COBOLT). We report one embedding for PBMC_1 (paired data) produced by COBOLT since it projects both modalities into a single embedding. Embeddings produced by scMMGAN appear inverted compared to the reference as it projects cells of one modality (*e.g.,* RNA) to the embedding learned from the other modality (*e.g.,* ATAC) and *vice versa*.

Figure S3: (A) Segmentation profiles of PDO used in the manuscript at 1 Mb resolution. Each row represents a single cell, segmentation values are colored from red (amplification) to blue (deletion). (B) Scatterplot representing the correlation between Chromatin Accessibility Score, calculated on scGET-seq data, and CRIS-B signature score, calculated on scRNA-seq data. Each point represents the median value of a score in cell groups at level 0 of the NSBM hierarchy. (C) Profile of gene signatures described for drug resistance and phenotypes and CRC subtypes. The heatmap reports the score of each signature in cell groups found in integrated data. (D) Profile of Adjusted Mutual Information (AMI) between PDO identity and cell groups identified using the Nested Stochastic Block Models. Orange dots indicate the groups from the integrated data, blue dots indicate groups identified in single modalities. The number near each dot indicates the level of the NSBM hierarchy. The red triangle indicates the level used for the analysis described in the manuscript.

Table S1: Performance of multiple tools in integration of PBMC data. Homogeneity, completeness and V-measure were evaluated between cell types and cell clusters. We report one value for COBOLT for each metric in PBMC_1 as it produces a single embedding for paired data. LISI scores were calculated on the input embedding (PCA) and the embedding generated by the tool. Both V-measure and LISI are reported in Figure 3 in the main text.

Table S2: Performance of integration of high number of modalities using MOWGAN. The table reports multiple scores calculated after four layer integration (RNA, ATAC, ADP and H3K27me3) of PBMC data, evaluated on the ability to transfer cell type annotation. The first three columns refer to label transfer from RNA to the remaining modalities. The other three columns report the values obtained by transferring labels from any other modality to RNA.

Table S3: Characterization of the PDO dataset. The table reports clinical features of three PDOs (CRC6, CRC17 and CRC39) and the mutational status of cancer genes commonly profiled in CRC. We also report the number of single cells analyzed in the corresponding scRNA-seq and scGET-seq datasets.

Table S4: Correlation between TF activity, computed on scRNA-seq data, and TBA scores, computed on scGET-seq data, in PDO datasets. We report Pearson’s *r* value and the associated *p*-value for transcription factors that were present in both analyses. Data are sorted by decreasing correlation.

