## Supplementary figures and images for "Scalable Integration of Multiomic Single Cell Data Using Generative Adversarial Networks"

### Supplementary Figure S1

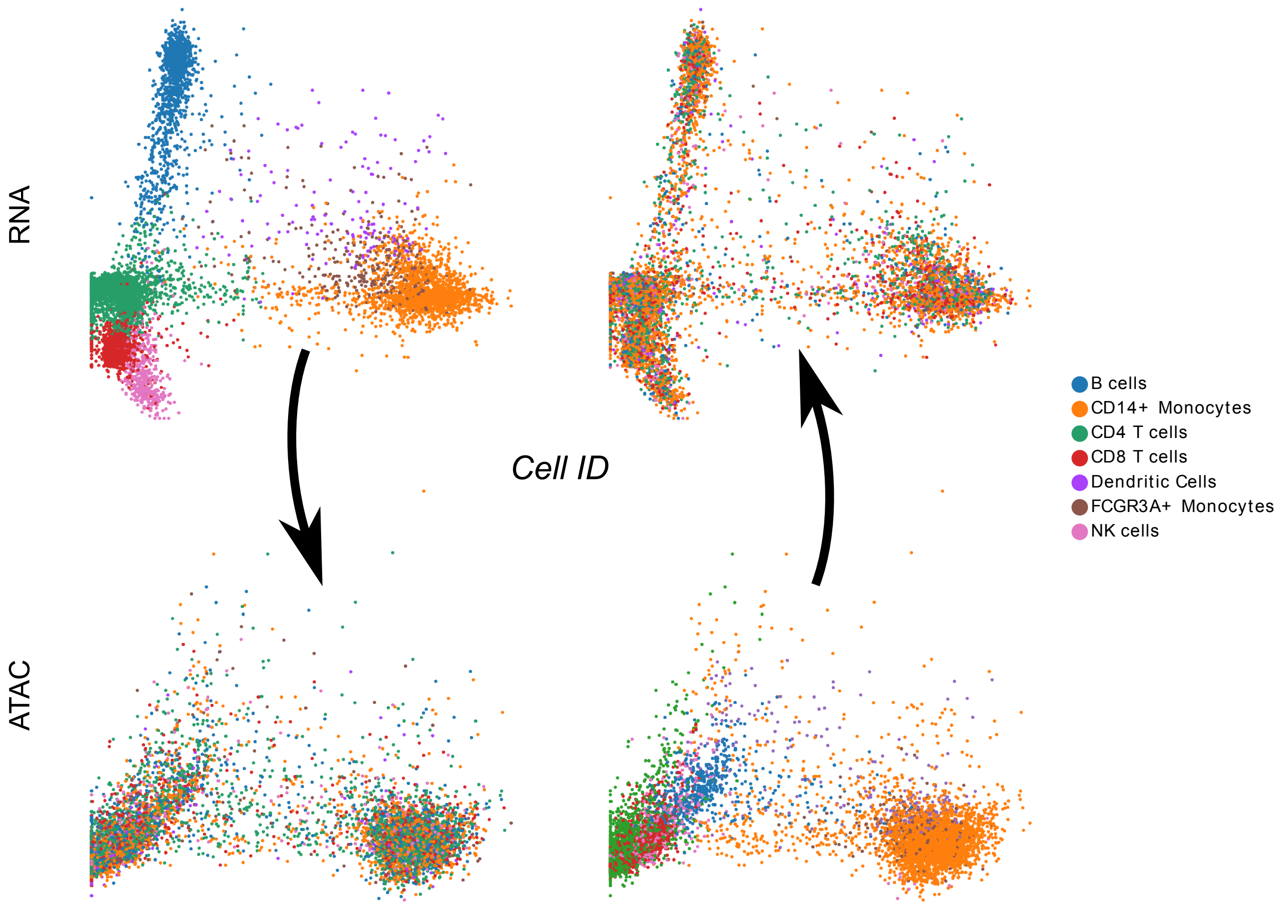

### Supplementary Figure S2

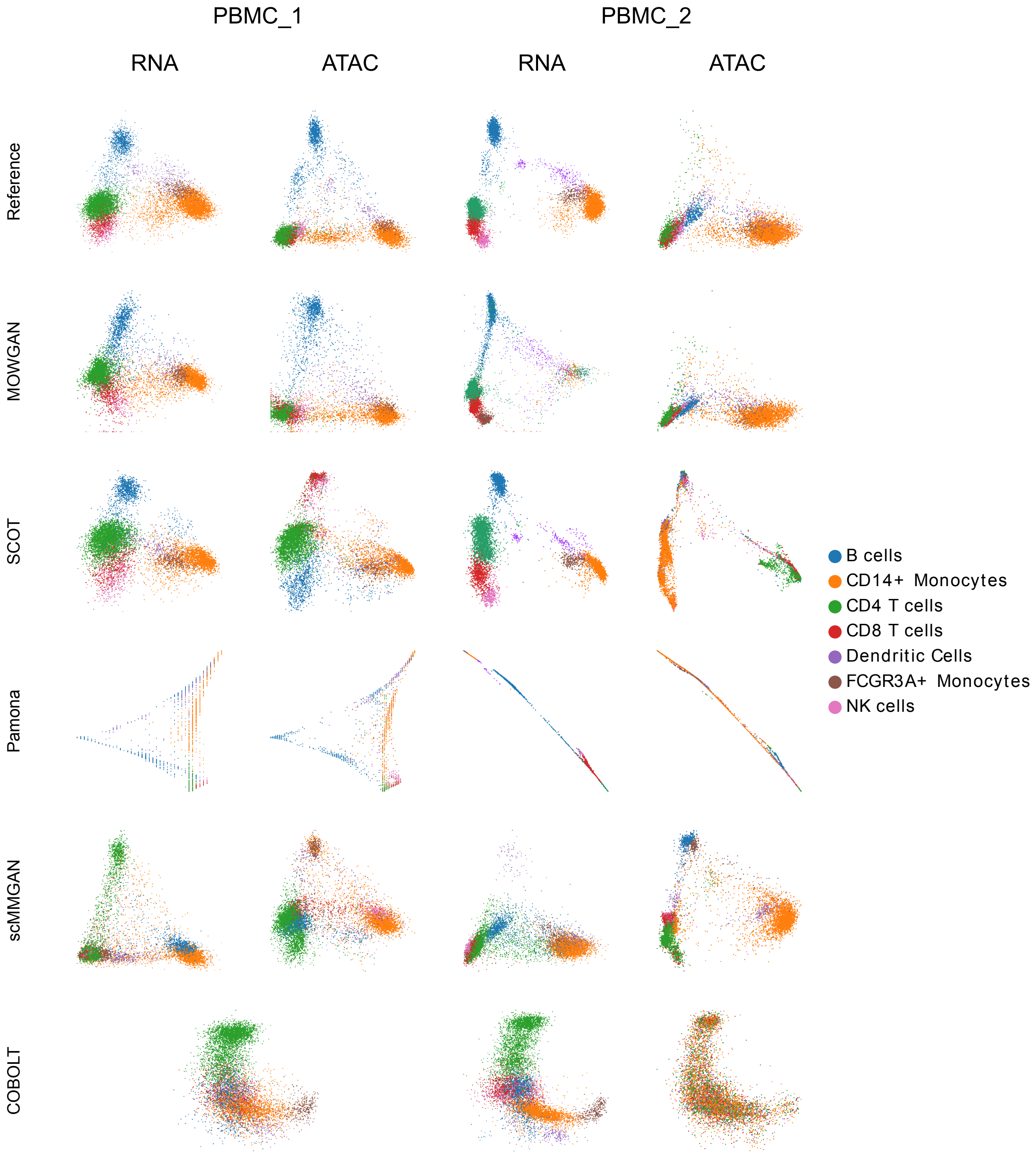

### Supplementary Figure S3

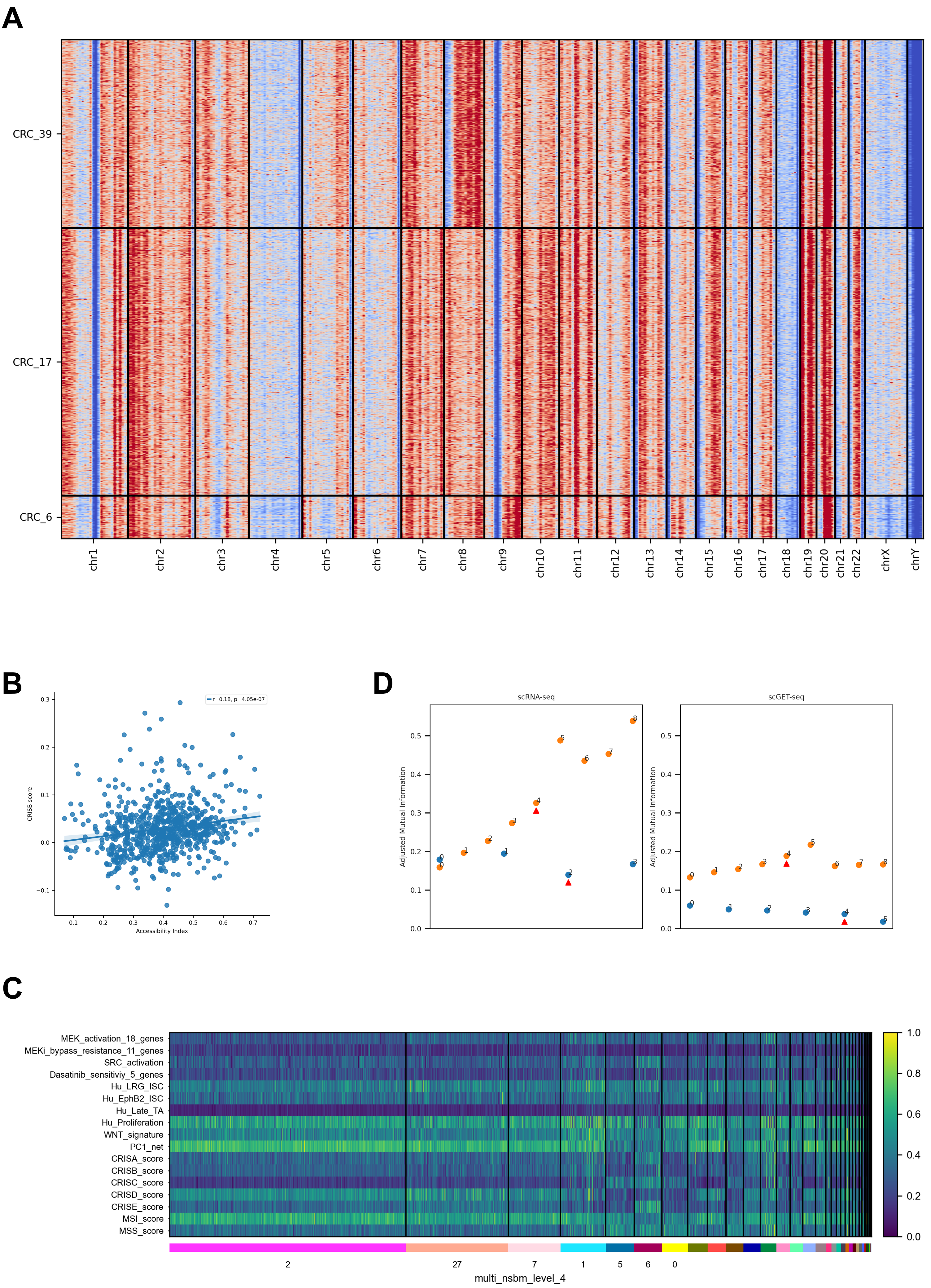
